# C19 trityl and silyl andrographolide analogs exhibit *in vitro* anticancer activity in murine cancer models

**DOI:** 10.64898/2026.10.27.762108

**Authors:** Rushika Raval, Ruirui Liu, Aileen Pak, Abigail Xinlan Yee, Srinidhi Venkatesh, Lutecia Lam, Alisha Hothi, Lavernie Chen, Allyson Yu, Hailey Seto, Claudia Casal Montserrat, Feng Wang-Johanning, Gary Johanning, Edward Njoo

## Abstract

Andrographolide, a labdane diterpenoid isolated from *A. paniculata* has been investigated for activity in a broad panel of biological contexts, including its anti-inflammatory, antiviral, and anticancer properties. Previously, A-ring analogs of andrographolide bearing silyl and trityl ethers at C19 have demonstrated improved *in vitro* anticancer activity in human cancer cell lines, with putative activity through the Wnt1/β-catenin signaling axis. In order to investigate the activity of this series of C19 andrographolide analogs, alongside a cyclic A-ring acetonide analog, as anticancer agents in murine cancer cell models, we evaluated antiproliferative activity of these compounds in B16F10 melanoma, CT26 colorectal adenocarcinoma, Neuro2a neuroblastoma, and 4T1 mammary ductal carcinoma cell lines, and found that most compounds exhibited potent micromolar level anticancer activity. Among these, all trityl ethers exhibited nearly indistinguishable IC_50_ values of 4-20 μM, and generally exhibited greater potency compared to C19 silyl ether analogs. This anticancer activity was further validated through apoptosis flow cytometry, wherein we concluded that our compounds induced apoptosis to a greater extent than the parent compound, andrographolide. Further, we examine the role of modulation of canonical Wnt1/β-catenin signaling in a series of co-delivery assays with CHIR-99021, a selective GSK-3β kinase inhibitor, and MSAB, a selective β-catenin degrader. Finally, we demonstrate that a selection of silyl ether analogs of andrographolide elicit cell cycle arrest by flow cytometry in a manner that is decoupled to *in vitro* potency trends. Altogether, we report the first *in vitro* evaluation of trityl and silyl ether analogs of andrographolide within murine adherent cancer cells.

## Introduction

Natural products and their analogs make up nearly a third of all small molecule therapeutics approved by the Food and Drug Administration (FDA) for the treatment of cancers^1–5^. Among naturally derived compounds, the vast structural diversity and architectural complexity of diterpenoids is accompanied by an even broader range of sophisticated biological activities and functions. Several examples of key diterpenoids approved by the Food and Drug Administration (FDA) for the treatment of human cancers include the taxanes paclitaxel, cabazitaxel, and docetaxel and ingenol mebutate, an ingenane ^6^. While natural products themselves continue to provide for the identification of small molecule candidates with unique anticancer potency, synthetic modifications have the potential to augment biological activities or alter endogenous functions of natural product leads ^7^.

An emergent approach in the design of new small molecules with improved potency or physiochemical properties leverages the functional repurposing of synthetic protecting groups such as silyl and trityl ethers ^8^. Our laboratory and others have demonstrated that the incorporation of organosilicon motifs on natural product scaffolds is a generalizable strategy for the modification of pharmacokinetic and pharmacodynamic properties of natural products, including Withaferin A ^9,10^, mycophenolic acid ^11^, lycorine ^12^, proscillaridin A ^13^, camptothecins, and the labdane diterpenoid andrographolide ^14–17^. Previously, Sirion et al. reported that the installation of silyl and trityl ether motifs at C19 of andrographolide greatly improved anticancer activity in P-388 (murine leukemia), KB (human oral epidermoid carcinoma), COL-2 (human colon cancer), MCF-7 (human breast cancer), LU-1 (human lung cancer), and ASK (rat glioma) cells ^17^. Subsequently, Reabroi et al. identified that a trityl ether analog of andrographolide exhibited single-digit micromolar potency in Wnt1-driven cancer cell lines through direct inhibition of nuclear localization and activation of β-catenin ^14,15^. More recently, our laboratory reported a systematic series of substituted silyl and trityl ether analogs of andrographolide ^16,18^ and found that these compounds exhibited improved potency compared to their parent natural product, with unique selectivity for HCT-116 and HT-29, two Wnt1-driven colorectal cancer cell lines. The role of Wnt1 inhibition was further previously established by a TCF/LEF Wnt1 reporter assay in this study.

Several groups have also reported the biological activity of the A-ring isopropylidene acetonide of andrographolide, which results in both increased A-ring rigidity and lipophilicity through functionalization of both C3 and C19 hydroxyl groups in a cyclic ketal. Previous studies have explored diverse aspects of the biological activity of this compound, including as an antiviral agent ^19,20^ and as an anticancer agent in murine P388 leukemia, human MCF-7 breast cancer, human actinic keratosis (AK), human SK-LU-1 (LU1) adenocarcinoma, and the human epithelial-like cancer cell line KB ^17^.

In several contexts, murine *in vitro* cancer models represent model proxies for human cancer counterparts, particularly in exploratory studies in fundamental signal pathway dependence^21,22^. Namely, several oncogenic pathways key to human tumor progression, including TCF/LEF dependent Wnt signaling, p53, and others, are highly conserved in murine models. Dysregulation of Wnt1/β-catenin signaling is also positively correlated with various murine carcinomas, including B16F10 murine melanoma, 4T1 mammary ductal adenocarcinoma, CT26 colorectal cancer, and Neuro2a glioblastoma cells. In CT26, a KRAS G12D mutant colon cancer cell line, Wnt10a, a primary ligand that activates the canonical Wnt1/β-catenin signaling pathway, is generally upregulated. Wnt1/β-catenin dependence is also observed in 4T1 and N2a cell lines, where the activation of this pathway aids in 4T1 tumor sphere formation and N2a cell-derived neuroblastoma formation ^23–25^. Alternatively, B16F10 tumor proliferation is associated with the upregulation of PI3K/AKT pathway, consequently leading to the phosphorylation of β-catenin ^26^. Moreover, some of these carcinomas overlap with the RAS signaling pathway, with N2a cells relying on downstream ERK/MAPK activation for neuronal differentiation and B16F10 and 4T1 exhibiting some dependence on RAS-mediated pathways such as NF-κB ^27^.

Collectively, the landscape of murine *in vitro* cancer models available and the potent anticancer activity of C19 andrographolide analogs previously reported in human cancer cell lines presents a unique opportunity to study the activity of these and similar compounds in murine models. Here, we expand our earlier report on the anticancer activity of C19 andrographolide analogs to four murine cancer models, and demonstrate that a significant degree of antiproliferative activity previously observed among andrographolide silyl and trityl analogs translates to murine models. We further evaluate an andrographolide A-ring acetonide, a model cyclic A-ring motif, which bears anticancer activity distinct from C19 silyl and trityl ether analogs **(Figure 1)**.

**Figure 1.**
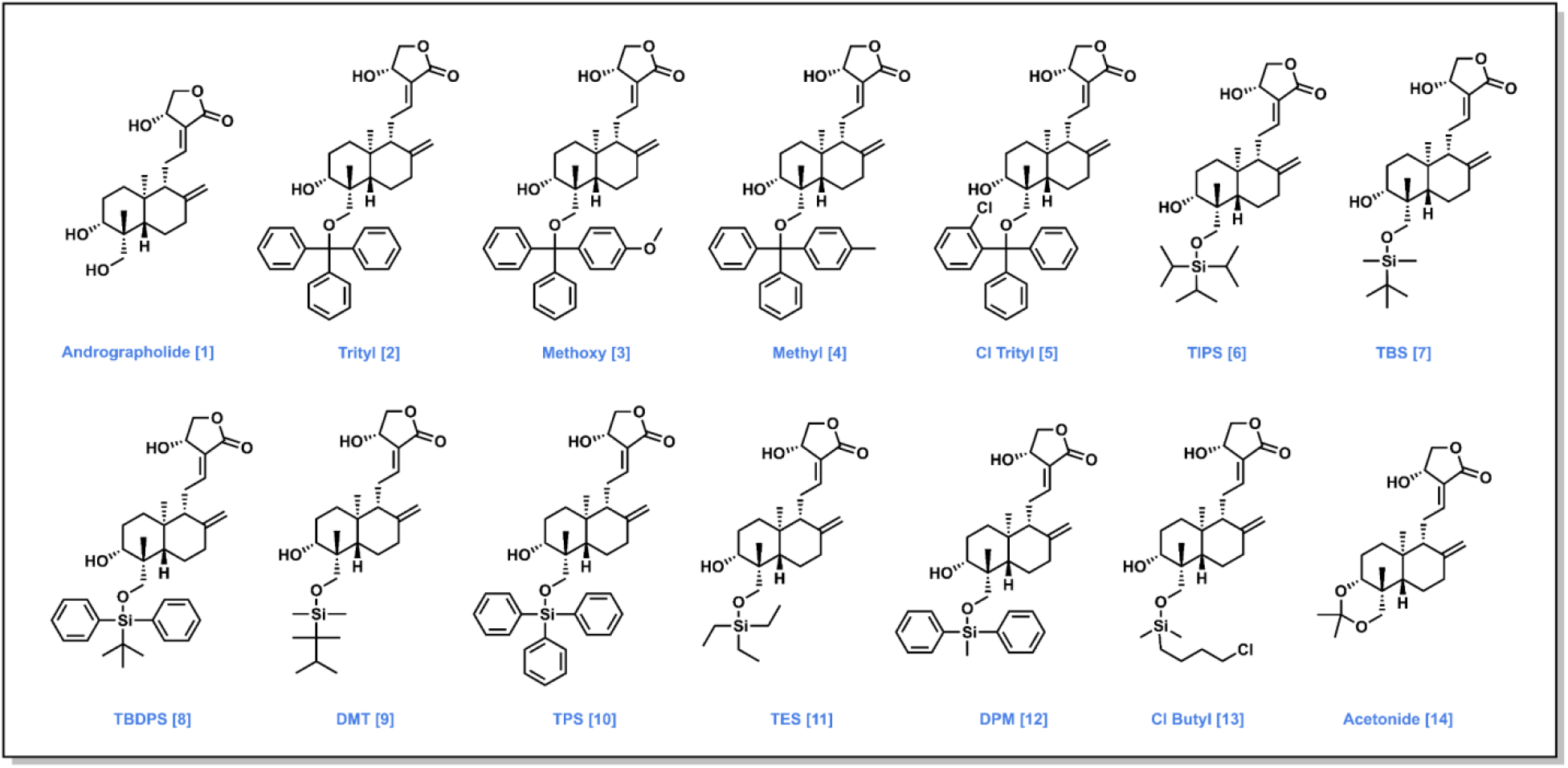
Library of andrographolide and analogs, **1-14**, featuring silyl and trityl ether substituents at C19 are evaluated for biological activity.

Across this panel of lipophilic A-ring functionalized andrographolide analogs, we observed potent antiproliferative activity in CT26, 4T1, B16F10, and Neuro2a cell lines, establishing the potential for these compounds in eliciting broad-spectrum *in vitro* anticancer activity in murine cancer models. Most compounds demonstrated exceptional potency in CT26 cells, with all compounds bearing sub-100 μM IC_50_ values. In order to determine if the observed anticancer activity of these compounds is mechanistically generalizable across other cancers, we estended this study across a panel of murine *in vitro* cancer models, including B16F10 murine melanoma, 4T1 mammary ductal adenocarcinoma, CT26 colorectal cancer, and Neuro2a glioblastoma cells. Given the augmented intracellular uptake of other silylated small molecules previously observed by our group ^28^, we further sought to identify trends of *in vitro* bioavailability through limited-exposure drug washout experiments and *in silico* evaluation of pharmacokinetic/pharmacodynamic parameters. In subsequent *in silico* PD/PK modeling of the physicochemical properties of these compounds, as well as drug washout experiments, silyl ethers and trityl ethers were found to exhibit moderately differentiated loss of potency when B16F10 or CT26 cells were subjected to briefer time points of drug exposure before washout, suggesting that silyl and trityl ether analogs of andrographolide demonstrate distinct kinetics of membrane-permeable bioavailability. We explored general mechanisms of antiproliferative activity of A-ring andrographolide analogs through both cell cycle and apoptosis flow cytometry. We further explored the potential role of Wnt1/β-catenin, as well as GSK-3β, through a series of co-delivery experiments and immunofluorescence imaging.

Beyond canonical Wnt1/β-catenin signaling, emerging evidence implicates human endogenous retrovirus type K (HERV-K) envelope (env) expression in the regulation of oncogenic signaling pathways associated with breast cancer progression ^29–31^. Previous studies have demonstrated elevated HERV-K env expression in human breast carcinomas and established functional relationships between HERV-K expression, tumor cell proliferation, invasion, and metastatic potential ^32^.

Mechanistically, modulation of HERV-K env expression has been associated with alterations in RAS/RAF/MEK/ERK signaling, while more recent studies have implicated additional changes in PI3K/AKT signaling, β-catenin expression, and epithelial-mesenchymal transition (EMT). These observations suggest that HERV-K-associated tumor biology may intersect with signaling pathways previously implicated in the anticancer activity of andrographolide and its C19-functionalized analogs. To investigate the activity of these compounds in a complementary tumor model, we employed a stably transfected 4T1-K murine mammary carcinoma cell line engineered to stably express full-length human HERV-K envelope.

Across these four murine cancer models, all C19 silyl and trityl ethers exhibit dose-dependent antiproliferative and pro-apoptotic activity, with similarly potent activity among substituted trityl ether analogs. Among silyl ethers, a dimethylthexylsilyl ether (Compound **9**) and triisopropylsilyl ether (Compound **6**) exhibited the greatest antiproliferative activity across cell lines, with IC_50_ values of 8.32 μM and 3.65 μM at 48 and 96 hours, respectively. Furthermore, washout experiments revealed even greater bioavailability differences between silyl and trityl ether analogs. Altogether, these results demonstrate that lipophilic functionalization at the C19 position of andrographolide generally leads to improved antiproliferative potency across B16F10, CT26, Neuro2a, and 4T1 murine cancer models, but that silyl and trityl ethers exhibit fundamental differences in washout experiments.

## 2. Materials and Methods

### 2.1 Chemical Synthesis

Compounds **2-14 (Figure 1)** were synthesized as previously described ^16^. Andrographolide acetonide was likewise prepared according to literature protocols. All compounds were characterized for identity and purity with ^1^H and ^13^C NMR, mass spectrometry, and Fourier-transform (FT-IR) spectroscopy.

### 2.2 Cell Culture

CT-26 murine colorectal cancer cell lines were cultured in RPMI-1640 media supplemented with 10% fetal bovine serum and 1% penicillin−streptomycin. 4T1, a murine breast ductal adenocarcinoma cell line, was cultured in RPMI-1640 medium, supplemented with 10% v/v fetal bovine serum and 1% v/v 100× penicillin−streptomycin. B16F10 murine melanoma cells were cultured in high glucose DMEM medium supplemented with 10% v/v fetal bovine serum and 1% v/v 100× penicillin−streptomycin. Neuro2a, a murine glioblastoma cell line, was cultured in EMEM media supplemented with 10% fetal bovine serum and 1% penicillin−streptomycin. All cells were grown to 80% to 100% confluency at 37 °C in a humidified incubator (5% CO_2_).

### 2.3 Cell Viability by MTT Assay

In order to determine dose-dependent and time-dependent antiproliferative activity of compounds **1-14 (Figure 2)** in murine cancer cell models, we performed a dose-titration colorimetric cell viability assay. Briefly, B16F10, CT26, 4T1, or Neuro2a cells were seeded at 30% confluency in tissue culture-treated 96 well plates. Following 24 hr incubation at 37°C / 5% CO_2_, compounds **1-14** were added as a 100× v/v DMSO dilution in DMEM, RPMI-1640, or EMEM media, respectively. Following an additional 48 hr or 96 hr incubation period, 10 μL of MTT solution (5.0 mg/mL in PBS) was added to each well and incubated with cells for 1 hour. Subsequently, the supernatant was aspirated off, and the resulting formazan crystals were redissolved in DMSO. Cell viability was quantified in a Molecular Devices M2 plate reader at 570 nm and normalized based on a 0.5% v/v DMSO control.

**Figure 2.**
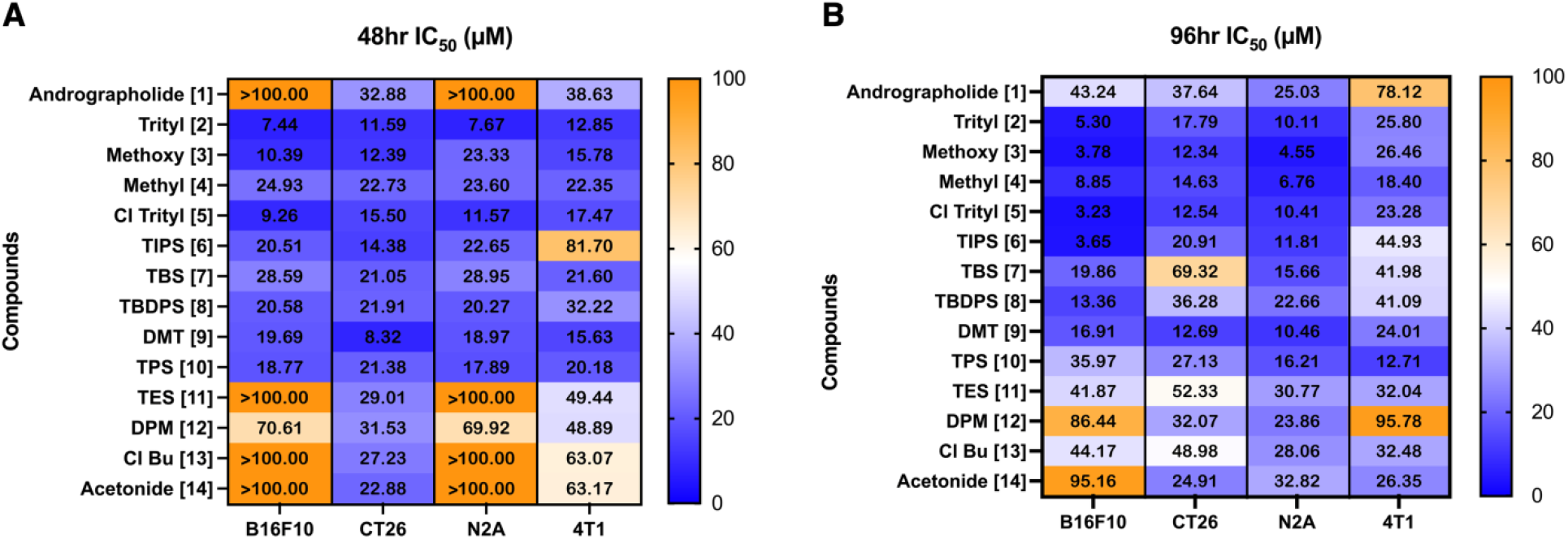
Antiproliferative Activity of andrographolide analogs in B16F10, CT26, neuro2A, and 4T1 murine tumor models. **A)** 48 hour IC_50_ values and **B)** 96 hour IC_50_ values, determined by MTT assay. Between cell lines, all compounds exhibit the most uniform potency at 48 hr in CT26 cells, while trityl ether analogs exhibit the greatest potency at 96 hr in B16F10 melanoma and Neuro2a glioblastoma cell lines.

### 2.4 Washout Experiments

In order to evaluate the effect of limited-time compound exposure on compound potency, we performed washout experiments in B16F10 murine melanoma and CT26 murine colorectal cancer cell lines **(Figure 3)**. Briefly, cells were seeded at 30% confluency in tissue culture-treated 96 well plates. Following 24 hr incubation at 37°C / 5% CO_2_, compounds **1-14** were added as a 200× v/v DMSO dilution in DMEM or RPMI-1640 media, respectively. After the cells were incubated with compound exposure for 2 hrs, the supernatant was aspirated off and displaced with fresh media. Following an additional 48 hr incubation period, cell viability was quantified via an MTT assay as previously described.

**Figure 3.**
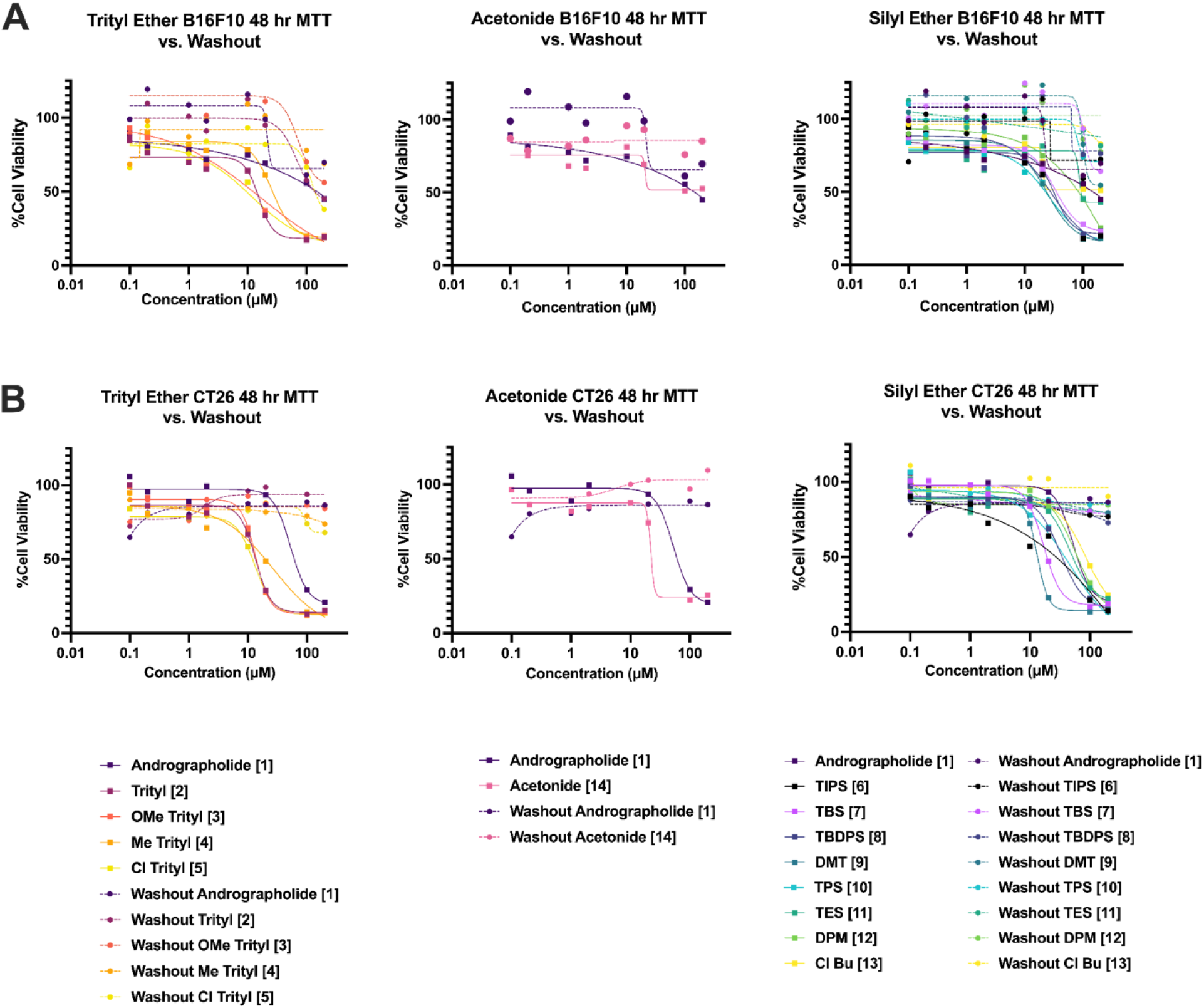
C19 silyl and trityl ether analogs exhibit decreased antiproliferative activity across **A)** B16F10 and **B)** CT26 cell lines following two hours of drug exposure and subsequent drug washout. Comparison of cell viability plots with and without drug washout for andrographolide and analogs **1-14** in B16F10 melanoma and CT26 colon carcinoma, respectively, quantified 48 hours after drug administration.

### 2.5 In silico PD/PK modeling

Chemical structures for compounds **1-14** were entered into SwissADME ^33^. Select physicochemical properties, including cLogP, MW, TPSA, LogS_w_, HBD+A, and RotBonds were plotted as a radar plot in Microsoft Excel **(Figure 4)**.

**Figure 4.**
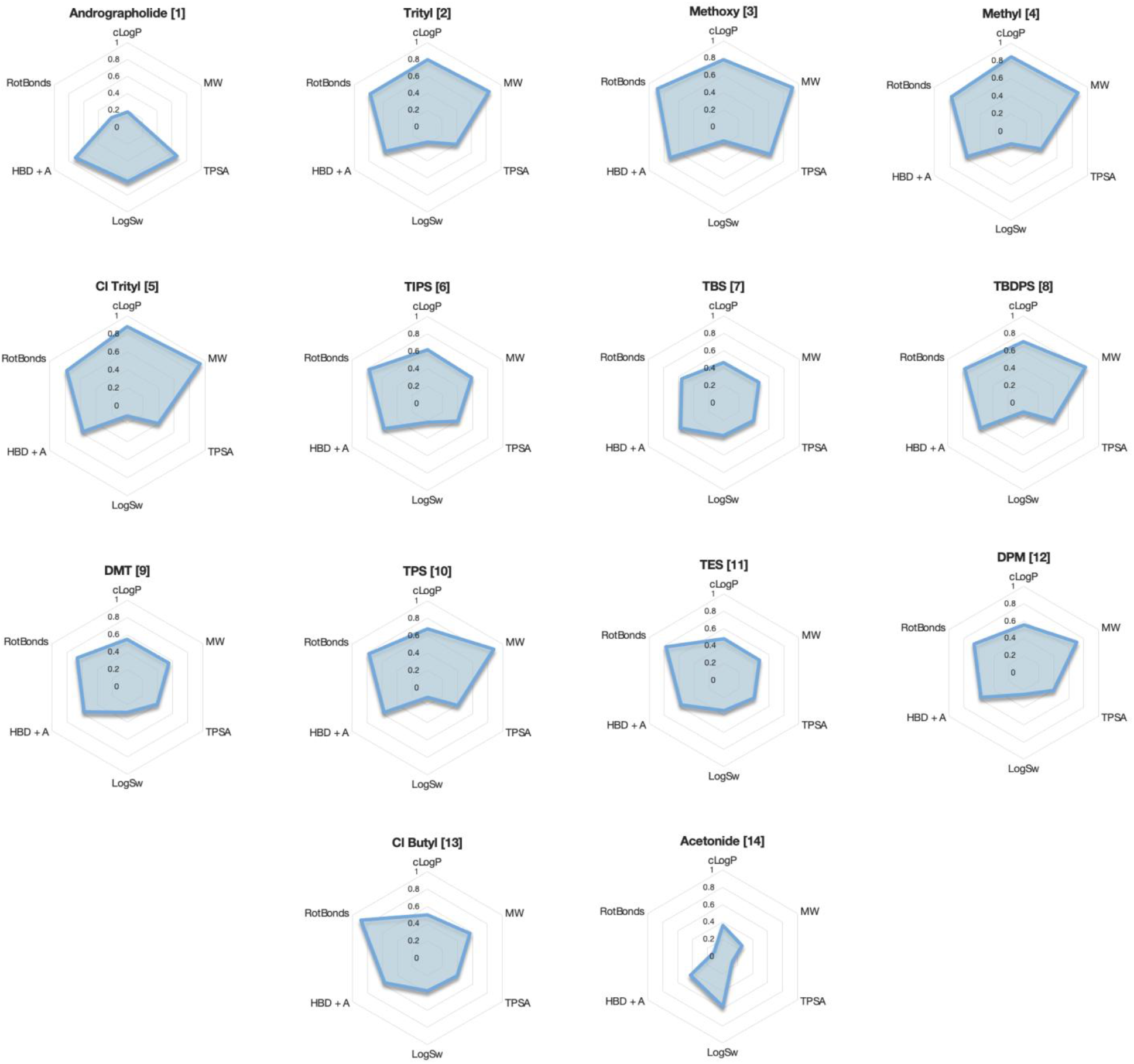
Predicted ADME properties of andrographolide and C19 functionalized analogs calculated by SwissADME are used to probe discrepancies in pharmacokinetic and pharmacodynamic properties for further insight into compound lipophilicity, solubility, and permeability in relation to experimental assessments of cytotoxicity. 1 ≤ cLogP ≤ 8; 300 ≤ MW ≤ 650; 60 Å^2^ ≤ TPSA ≤ 100 Å^2^; −10 ≤ LogS_w_ ≤ 0; 3 ≤ HBD + A ≤ 10; 1 ≤ RotBonds ≤ 10.

### 2.6 Co-delivery with CHIR-99021

In order to evaluate potential GSK-3β dependent Wnt1 activity, we performed a series of co-delivery experiments of C19 andrographolide analogs and CHIR-99021, a previously reported small molecule GSK-3β inhibitor, in CT26, B16F10, and 4T1 cells **(Figure 5)**. Briefly, cells were seeded in 96 well clear flat bottom tissue culture plates. Following 24 hr incubation at 37°C / 5% CO_2_, compounds **1-14** were added as a 100× v/v DMSO dilution in CHIR-99021 laced DMEM, RPMI-1640, or EMEM media, respectively, at 10 µM final concentration. Following an additional 48 hr or 96 hr incubation period, 10 μL of MTT solution (5.0 mg/mL in PBS) was added to each well and incubated with cells for 1 hour. Subsequently, the supernatant was aspirated off, and the resulting formazan crystals were re-dissolved in DMSO. Cell viability was quantified in a Molecular Devices M2 plate reader at 570 nm and normalized based on a 0.5% v/v DMSO control.

**Figure 5.**
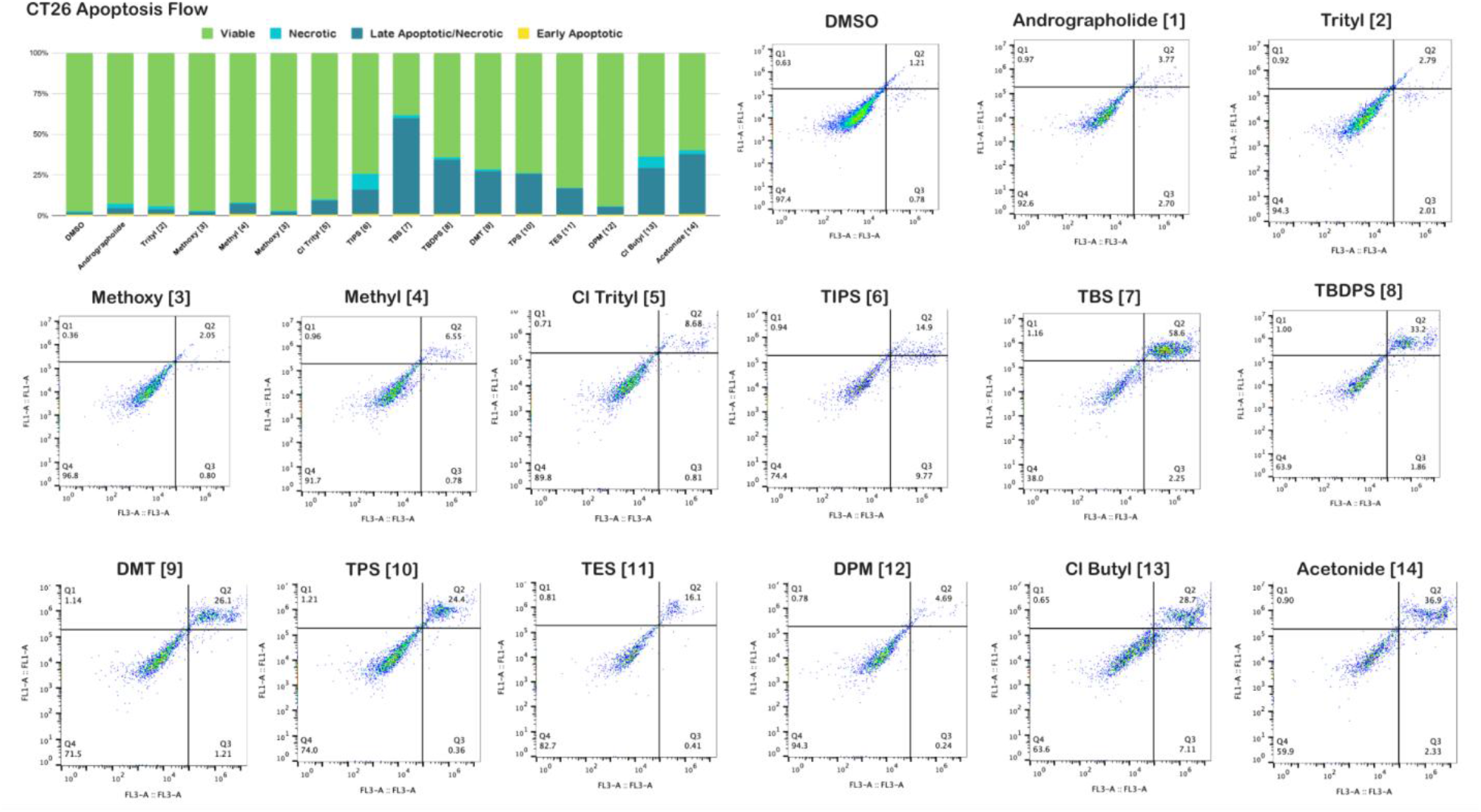
Select C19 silyl analogs demonstrate an increased late apoptotic/necrotic cell population percentage in CT26 colon carcinoma through apoptosis flow cytometry analysis. In cells treated with analogs **1-14** and a DMSO negative control for 24 hours, cells were labeled with FITC-Annexin V and propidium iodide (PI) and were measured by flow cytometry for apoptotic or pro-apoptotic activity.

### 2.7 Apoptosis Flow Cytometry

Cell cultures were prepared referencing the protocol above. Briefly, CT-26 cells were seeded at 40,000 cells/mL per well in 6 well flat-bottom tissue culture-treated plates and incubated in a humidified incubator for 24 hrs at 37°C / 5% CO_2_. Following the 24 hr incubation period, the cells were treated with compounds **1-14** at a desired concentration of 500 nM. A negative control was established by treating cells with 0.5% v/v DMSO. After incubation for 24 hrs, the cell medium was aspirated off into separate Falcon tubes; the cells were trypsinized with 500 μL of 0.25% trypsin−EDTA and incubated for 15 min. After quenching with 1 mL of cell-line specific media, detached cells were aspirated off into the Eppendorf tubes and centrifuged until cell pellet formation. The supernatant medium was removed, and the cell pellet was resuspended in 100 μL of 1× PBS before additional centrifugation. The PBS was aspirated off, and the pellet was resuspended with 100 μL of 1× Annexin-V FITC Binding Buffer before being transferred to 1 mL Eppendorf tubes. The cells were then co-stained in the dark for 15 min at room temperature with 5 μL of Annexin-V FITC and 1 μL of propidium iodide stain (1 mg/mL in DI water, AK Scientific, 98%). The flow cytometer readings were obtained with a BD C6 Accuri Flow Cytometer and processed with FlowJo Software 10 using wavelengths FL-1A 533/30 nm and FL-3A > 670 nm filters.

### 2.8 Cell Cycle Flow Cytometry

Cell cultures were prepared referencing the protocol above. CT-26 cells were seeded at 80% confluency and incubated in a humidified incubator at 37°C / 5% CO_2_ for 24 h. Following 24 hrs of incubation, the cells were treated with compounds **2, 3, 5, 7, 8, 10, and 14** at 50 μM. A negative control of 0.5% v/v DMSO was also included. After incubation for an additional 24 hrs, the supernatant was aspirated off into Eppendorf tubes, and cells were trypsinized with 500 μL of 0.25% trypsin−EDTA per well and incubated for 10 min. After quenching with 500 μL of cell-line specific media, detached cells were moved into their respective Eppendorf tubes and centrifuged for 5 min at 2500 rpm until a cell pellet formed. The supernatant medium was removed, and the cell pellet was resuspended in 100 μL of 1× PBS and transferred into a 1 mL Eppendorf tube. After centrifuging for an additional 4 min at 2500 rpm, the PBS was removed, and cells were fixed in 500 μL of cold 70% ethanol for 30 min before centrifuging for an additional 4 min. The ethanol was aspirated off, and cells were resuspended in 200 μL of PBS and 50 μL of RNase (1 mg/mL, RPI), followed by incubation for 5 min at 37°C. The cells were then stained in the dark with 1 μL of propidium iodide (PI) stain for 30 min. Cell cycle analysis was run on a BD C6 Accuri Flow Cytometer, and analysis of results was performed through BD C6 Accuri Software using the FL-2A 585/40 nm filter. Results were processed using FlowJo Software 10 based on PI/count gating to determine the percentage of cells in the G0/G1 phase, S phase, and G2/M phase of the cell cycle for each sample.

### 2.9 Immunofluorescence Imaging

B16F10 or CT26 cells were seeded in glass-bottom 96-well plates and incubated for 24 hours at 37°C / 5% CO_2_. Afterwards, a drug medium was created by dissolving 2 mM solutions of compounds **1-14** in DMSO, which were then added to media to reach the desired concentration of 10 μM. Two columns were used for negative controls with cells and media only and with 0.5 % v/v DMSO. After incubation for 24 hours, the media was removed and the cells were fixed with the addition of methanol (Acros Organics) and incubation at 37°C for 20 minutes. Subsequently, 50 μL of 1.0% Triton X (BioPharm Inc.) was added and the cells were incubated at 37°C for another 30 minutes. The lysis buffer was removed and the cells were immediately washed with PBS and covered with aluminum foil before being stored in the refrigerator overnight at 4°C. The following day, two solutions were diluted 200-fold with 1.0% bovine serum albumin (BSA) solution from a solution of mouse anti-active-β-catenin primary monoclonal antibody (MilliporeSigma #05-665). 20 μL of the anti-active-β-catenin antibody solution was added to each well. After storage overnight at 4°C, the antibody solutions were removed and the cells were washed with PBS. Goat anti-mouse IgG(H+L)-AF488 antibody (Southern BioTech, #1036-30) was diluted 200-fold in 1.0% BSA solution before addition of 20 μL of the solution to each well. After incubation in the dark at 4°C for 3 hrs, the secondary antibody solution was aspirated off, and the cells were washed twice with 50 μL PBS. Then, a 14.3 mM solution of DAPI in DMF was diluted 50-fold in PBS, before another dilution of 1000-fold in PBS, to result in a final concentration of 300 nM. After washing with PBS twice, 20 μL of this solution was added to each well for 5 minutes at 4°C before being immediately washed with PBS twice.

Cells were imaged with a Zeiss Axiovert 200 Fluorescence Microscope with an Objective LD Achroplan 32x/0.40 Korr Ph2 lens. Overlay and final images were processed using ImageJ according to what has been previously described ^14,16^. Additionally, the total fluorescence at 485 nm was measured on a plate reader (Molecular Devices, Paradigm)

To quantify nuclear-associated β-catenin fluorescence, DAPI-channel images were used to generate binary masks delineating nuclear regions. Corresponding AF488-channel images were converted to grayscale, and the fluorescence intensity of pixels spatially overlapping with the binary nuclear masks was quantified. The resulting nuclear-associated fluorescence intensity, expressed as a percentage of the maximum possible grayscale pixel intensity, was used as a proxy for nuclear β-catenin accumulation. Images were randomly segmented into four partitions, and the resulting percent nuclear β-catenin accumulation is graphed with an average and standard deviation of the four independent calculations.

### 2.10 Generation of stably-transfected HERV-K 4T1 cell line

A full-length HERV-K *env* sequence from a viral particle isolated from a BC patient diagnosed with invasive ductal carcinoma was cloned into vector pLVX-DsRed-Monomer-C1 to construct the HERV-K *env*-expressing vector pLVX-Kenv. The HERV-K *env* expressing lentiviral particles were then packaged and titered in 293FT cells according to the manufacturer’s instructions (Lentivector Expression Systems: SBI System Biosciences), except using Lipofectamine 2000 instead of Fugene. The empty vector pLVX-DsRed-Monomer-C1 was also packaged to form the control lentiviral particles (pLVX or pLVXC).

### 2.11 In vivo models

BALB/c mice were procured from Jackson ImmunoResearch Labs. 4T1-K cells, stably transfected as previously described, were injected subcutaneously into the mice at 5-6 weeks of age (2 x 10^6^ cells). Five days after cellular inoculation, a dosing regimen of 0.5 mg/kg andrographolide (Compound 1), *tert-*butyldiphenylsilyl ether (Compound 8), dimethylthexyl silyl ether (Compound 9), or DMSO vehicle control (10% v/v in 5% saline, prepared in 200 μL injections) was administered QOD twice weekly via intraperitoneal injection for twenty-seven days. Tumor sizes and body weights were recorded every 48 hours. Twenty-eight days after cellular inoculation, all mice were euthanized and endpoint tumor weight as well as tumor dimensions were measured. Volume was calculated by length (L) and width (W) using the formula (volume = π/6 x L x W^2^). A low-flow digital anesthesia delivery system (The Kent Scientific SomnoSuite) was employed to deliver controlled percentages of inhalant anesthetics, like isoflurane, following the manufacturer’s recommendations, to achieve a loss of the righting reflex. We performed cervical dislocation on the fully anesthetized, unresponsive animal to ensure completeness. All animal experiments were approved by the IACUC of The University MD Anderson Cancer Center IACUC (protocol no. 01-04-00733), SRI International (protocol no. 14001), and SunnyBay Biotech (protocol no. SBB-2020-01).

## 3. Results & Discussion

### 3.1 Anticancer Activity of C19 Andrographolide Analogs

Across all four murine cancer cell lines, both classes of C19-functionalized analogs exhibited increased antiproliferative potency compared to the parent compound andrographolide. Between silyl and trityl ether analogs, trityl ethers **2-5** exhibited marginally more potent antiproliferative activity. This trend appears to be consistent across all four cell lines, with limited time-dependence of IC_50_ values in 48 hr **(Figure 2A)** and 96 hr exposure runs **(Figure 2B)**. Acetonide **14**, which bears an A-ring cyclic ketal in lieu of C19 silyl or trityl functionalization, exhibits a time-dependent decrease in IC_50_ in Neuro2a cells and unique potency for CT26 cells at 48 hr timepoint in a manner similar to andrographolide. Among silyl ethers, dimethylthexylsilyl ether **9**, and to some extent, triphenylsilyl ether **10**, exhibited marginally higher potency than other silyl ether analogs. Like the trityl ether analogs, this class of C19 silyl ether analogs did not exhibit significant IC_50_ differences at 48 hrs and 96 hrs of exposure, suggesting that the primary mechanism through which these compounds operate experiences saturation of dose-dependency before 48 hrs of exposure.

### 3.2 Washout Experiments

To assess the cellular uptake of these silyl and trityl ether analogs, we conducted washout assays in B16F10 cells **(Figure 3A)** and CT26 **(Figure 3B)**. After two hours of exposure to compound-laced media, the residual supernatant was removed and replaced with new, undrugged media in order to measure the relative potency of the partition of compound dosing that was internalized intracellularly after only two hours of compound exposure. Expectedly, all compounds exhibit a precipitous loss of potency during this limited-exposure experiment, suggesting that silyl and trityl ether derivatives of andrographolide are only partially internalized into cells over the course of the two hour exposure time **(Figure 3)**. Between silyl and trityl ether analogs, all trityl ether analogs exhibit similar losses in potency. Additionally, the acetonide (Compound **14**) exhibits a near total loss of antiproliferative potency in CT26 cell lines.

From this assay, there also appears to be slight differences in resultant potency between washout assays conducted in B16F10 cells as compared to CT26 cells; in B16F10 cells, silyl and trityl ethers appear to retain a slightly greater potency after only two hours of exposure as compared to CT26 cells, in which nearly all compounds experience a total loss of biological potency. This suggests that the rate of membrane penetration and relative bioavailability baselines may be unique to different cell lines.

### 3.3 In silico modeling

*In silico* calculated physicochemical properties of andrographolide and its analogs are graphed on a radar plot **(Figure 4)**. Expectedly, all A-ring functionalized analogs exhibit greater lipophilicity, as evidenced by much higher cLogP values compared to andrographolide (2.26) **(Table 1)**. Generally, trityl ethers **2-5** exhibited higher cLogP values (6.48-7.15) compared to their silyl ether counterparts **6-13**, which exhibited cLogP values ranging from (4.23-5.91). Expectedly, both silyl and trityl ethers exhibited greater cLogP values compared to the A-ring acetonide (3.36).

**Table 1.**
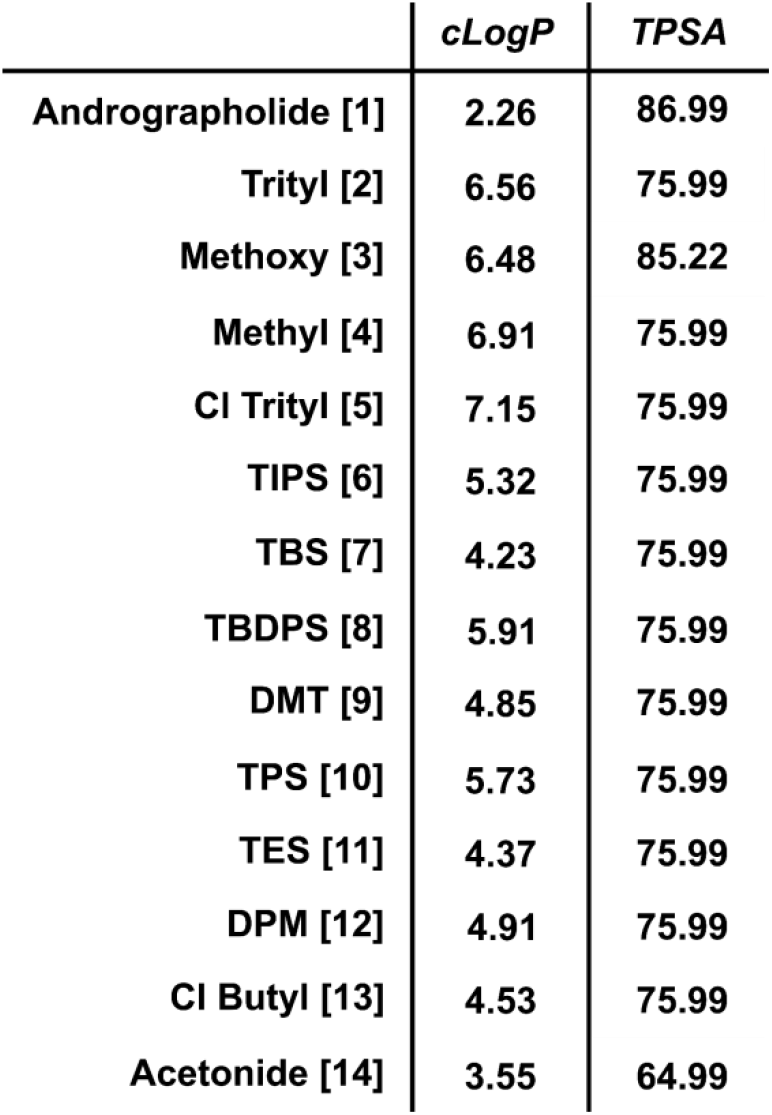
Predicted cLogP and TPSA for andrographolide and A-ring functionalized analogs indicate an overall greater lipophilicity of A-ring analogs as compared to andrographolide, with trityl ethers exhibiting higher cLogP values than the silyl ethers. Both silyl and trityl ethers display greater lipophilicity than the andrographolide acetonide, while predicted TPSA values remained relatively uniform across all compounds.

|  | *cLogP* | *TPSA* |
| --- | --- | --- |
| Andrographolide [1] | 2.26 | 86.99 |
| Trityl [2] | 6.56 | 75.99 |
| Methoxy [3] | 6.48 | 85.22 |
| Methyl [4] | 6.91 | 75.99 |
| Cl Trityl [5] | 7.15 | 75.99 |
| TIPS [6] | 5.32 | 75.99 |
| TBS [7] | 4.23 | 75.99 |
| TBDPS [8] | 5.91 | 75.99 |
| DMT [9] | 4.85 | 75.99 |
| TPS [10] | 5.73 | 75.99 |
| TES [11] | 4.37 | 75.99 |
| DPM [12] | 4.91 | 75.99 |
| Cl Butyl [13] | 4.53 | 75.99 |
| Acetonide [14] | 3.55 | 64.99 |

Also consistent with our observations, all silyl ethers and trityl ethers of andrographolide exhibited much lower aqueous solubility values, as demonstrated by significant decreases in LogS_w_ values ranging from −8.95 to −6.20 for silyl ethers, and −8.88 to −8.22 for trityl ethers. All compounds had a far greater molecular weight compared to andrographolide, approaching and in some cases exceeding Lipinski’s 500 MW ceiling.

### 3.4 Apoptosis cell cycle flow cytometry

Given the unique potency of these compounds observed in CT26 cells, we selected this cell line for quantification of apoptotic activity by flow cytometry. Expectedly, cells treated with a DMSO control exhibited minimal levels of apoptotic activity, while the administration of several silyl ethers at 50 μM resulted in a significant increase in apoptotic cells. Notably, cells treated with *tert*-butyldimethylsilyl analog **7** exhibited the greatest pro-apoptotic activity (59.76%), while the A-ring acetonide **14** exhibited comparable pro-apoptotic activity as compared to *tert-*butyldiphenylsilyl ether **8** and 4-chlorobutyldimethyl ether **13 (**37.8%, 34.2%, 29.35%, respectively) **(Figure 5)**.

Interestingly, while *in vitro* cell viability experiments indicate potent antiproliferative activity across all cell lines elicited by trityl ethers **2**-**5**, these compounds resulted in minimal induction of pro-apoptotic activity. It is also noteworthy that triphenylsilyl ether **10** exhibits very different pro-apoptotic activity in CT26 cells as compared to trityl ether **2** (25.61% and 3.71% apoptotic, respectively), despite only differing in a single atom-analogous carbon-to-silicon substitution. This suggests that C19 trityl ethers of andrographolide, despite having similar antiproliferative potencies as C19 silyl ethers, operate through a distinct mechanism that may not generate as profound a pro-apoptotic effect measured by FITC-Annexin V.

### 3.5 Cell Cycle Flow Cytometry

We additionally probed whether the addition of andrographolide analogs results in changes to cell cycle distribution in CT26 cells. We observed no correlation between the selective pro-apoptotic activity elicited by silyl ethers **6-13** and unique changes in cell cycle distribution between G1, S, and G2/M phase, with all trityl ethers and most silyl ethers exhibiting very similar cell cycle distributions as the DMSO control **(Figure 6)**. Administration of triphenylsilyl ether **10** resulted in moderate G2/M phase arrest. Similarly, in 4T1 cells, minimal changes to cell cycle distribution were observed across all compounds. Collectively, these results suggest that cell cycle arrest is not a primary mechanism of action of andrographolide C19 analogs in CT26 murine colorectal cancer cells.

**Figure 6.**
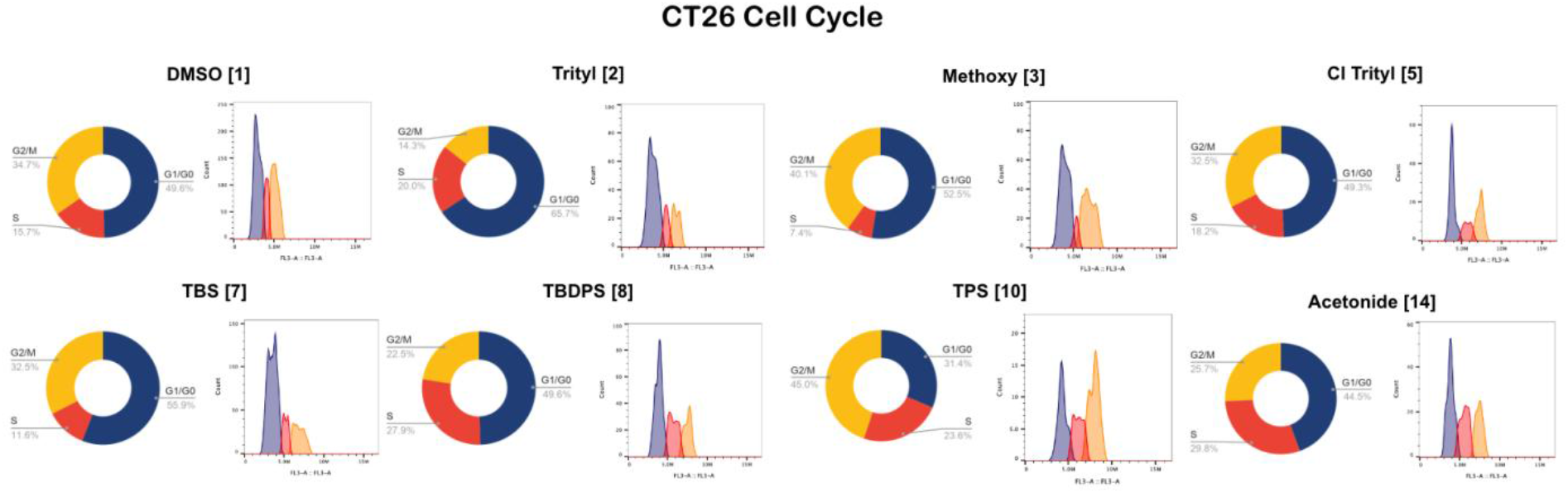
Cell cycle flow cytometry analysis of andrographolide and C19 analogs in CT26 colon carcinoma cells exhibits minimal cell cycle arrest.

### 3.6 Co-delivery with CHIR-99021, a GSK-3β Inhibitor

Next, to investigate the potential role of Wnt signaling and GSK-3β, which we and others have previously explored as putative mechanisms through which andrographolide C19 analogs exhibit anticancer activity in human cancer cell lines, we evaluated changes in antiproliferative IC_50_ values when compounds **1-14** were administered in combination with 10 μM CHIR-99021.

In 4T1 murine breast duct adenocarcinoma cells and B16F10 melanoma cells, coadministration with CHIR-99021 did not result in significant differences in potency **(Figure 7)**, with IC_50_ values within experimental range of cells treated with most compounds in this series. Moderate loss of potency was observed when 4T1 and B16F10 cells were treated with andrographolide (Compound **1)**, triethylsilyl ether (Compound **11)**, diphenylmethylsilyl ether (Compound **12)**, chlorobutyldimethylsilyl ether (Compound **13**) and the A-ring acetonide (Compound **14)**. Remarkably, compound **14** results in the greatest loss of potency upon codelivery with 10 μM CHIR-99021, suggesting that this compound may act through greater GSK-3β dependence than silyl and trityl ethers of andrographolide.

**Figure 7.**
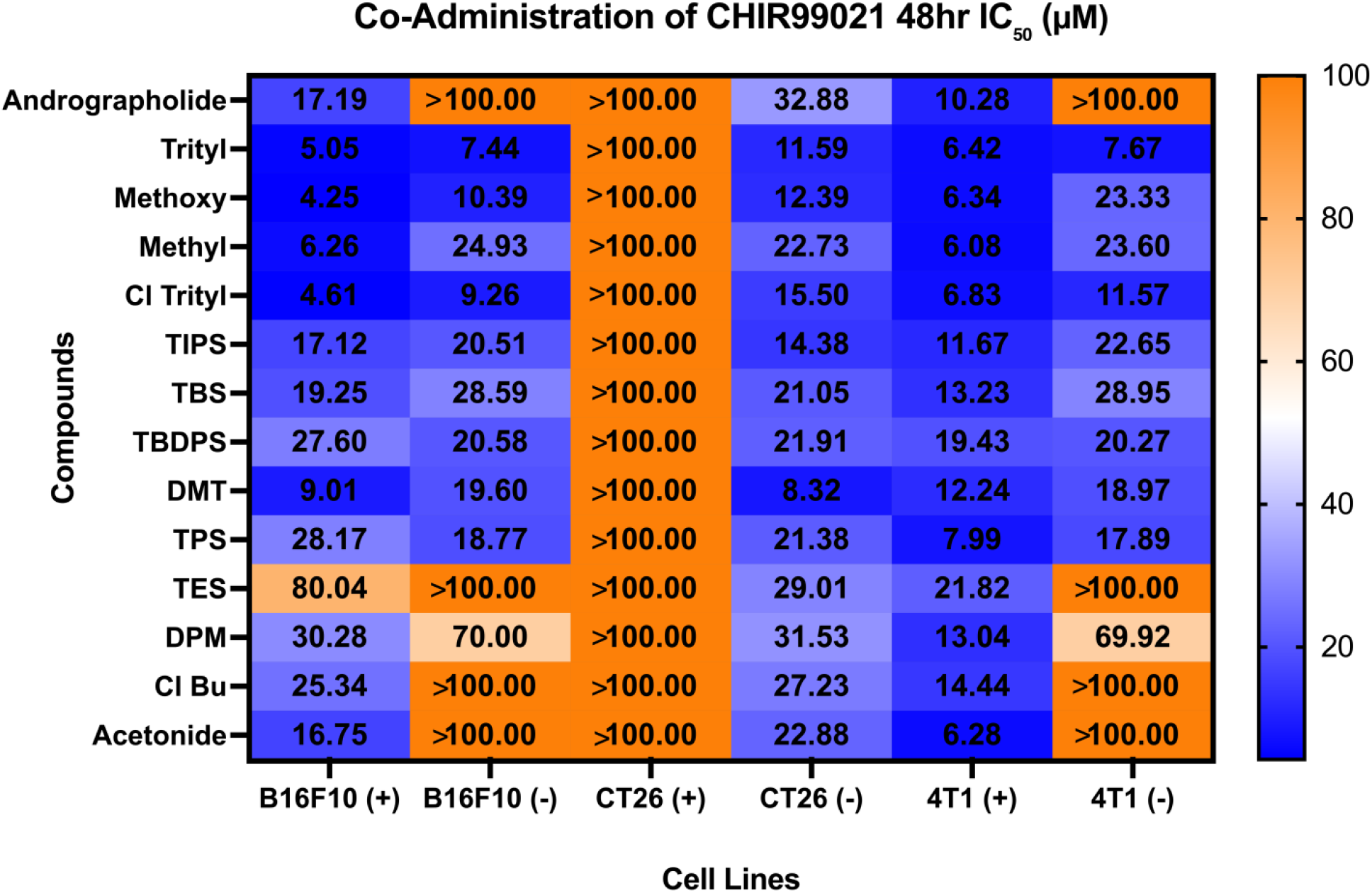
Co-administration of GSK-3β inhibitor CHIR-99021 (10 µM) in B16F10 melanoma, 4T1 mammary carcinoma, and CT26 colorectal cancer cells results in different degrees of changes in antiproliferative potency after 48 hours of drug administration, suggesting that different degrees of GSK-3β modulation of β-catenin is implicated in the compounds’ primary mechanism of action.

However, distinct from these two cell lines, the co-administration of CHIR-99021 in CT-26 resulted in a precipitous loss of potency, suggesting that the andrographolide A-ring analogs explored here exhibit unique sensitivity to GSK-3β-dependent Wnt signaling that is specific to CT26 murine colorectal cancer cells.

### 3.7 Immunofluorescence Imaging

Next, to visualize the cellular distribution of β-catenin, which is the putative downstream effector of the Wnt1 signaling axis whose nuclear localization was previously reported to be targeted by C19-trityl andrographolide ^14^, we conducted immunofluorescence imaging where cells treated with each compound were fluorescently stained for β-catenin and counterstained with DAPI. The resulting plates were imaged at 488 nm and 405 nm to determine intracellular distribution of β-catenin in both cytosol and nucleus. We further measured concentration-dependent changes in intracellular total β-catenin with a fluorescence plate reader. In both visual quantification from imaging experiments and direct fluorescence measurements, we found that there are no significant changes to total β-catenin content in cells treated with C19 andrographolide analogs, consistent with previous studies **(Figure 8)**. In B16F10 cells treated with 5 μM of compounds **1 – 14,** significant cell rounding was observed after 24 hours of exposure; CT26 cells were unable to be imaged at this concentration due to significant loss of cell viability within 24 hours, and thus were imaged at 1 μM final concentration.

**Figure 8.**
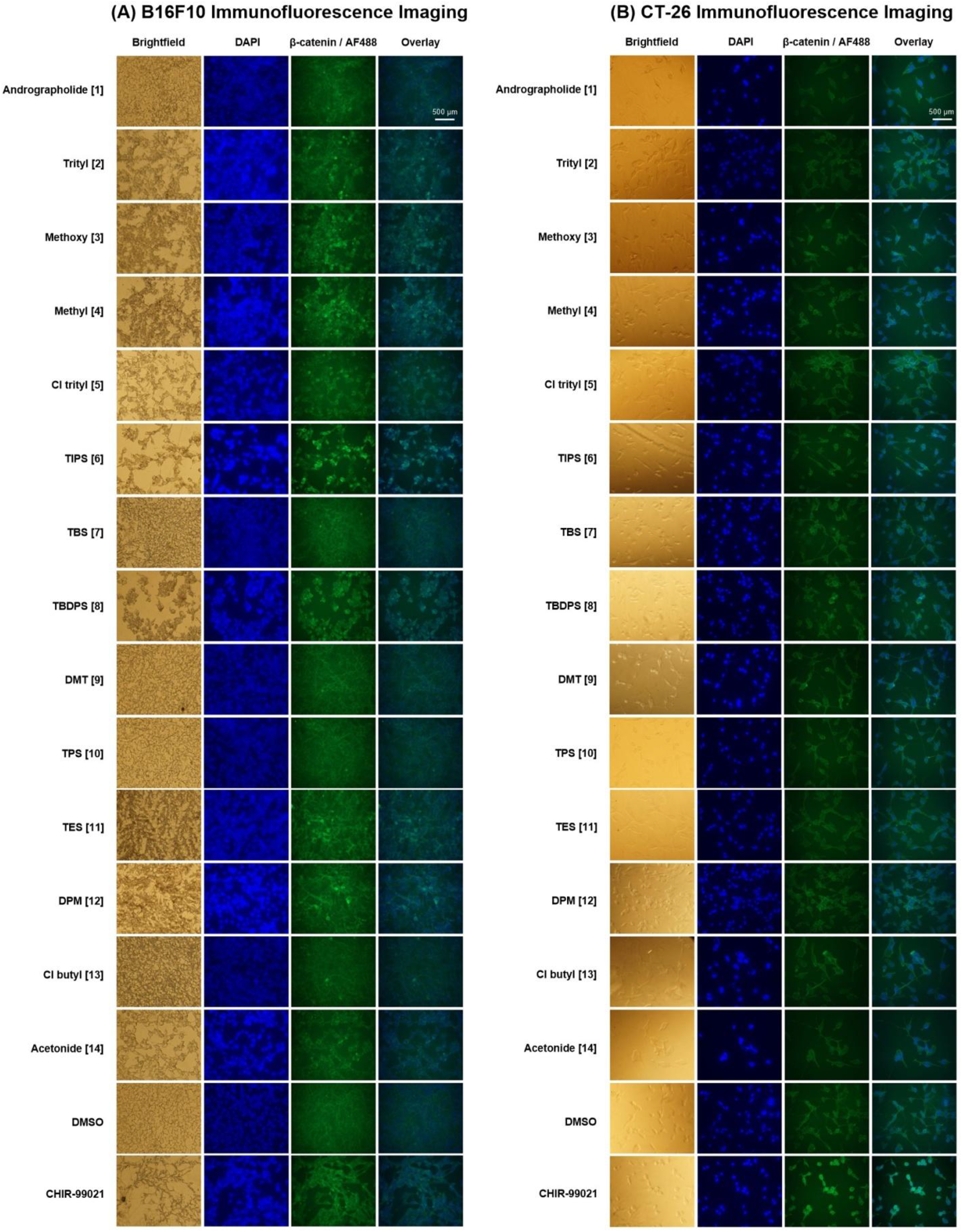
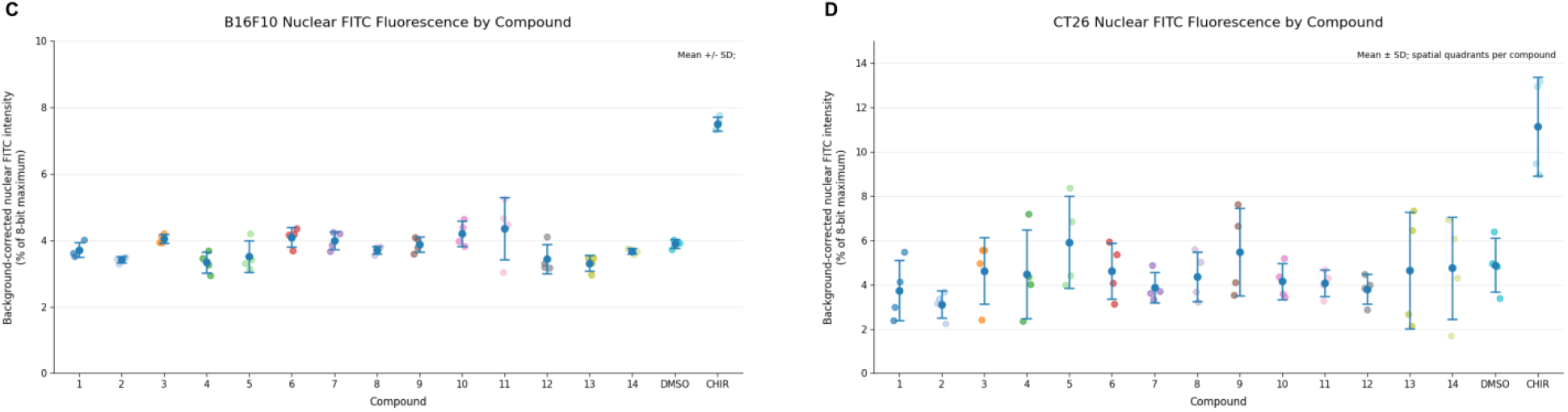
Immunofluorescence imaging of β-catenin in murine cancer cells. **A)** B16F10 cells were fixed and stained for beta catenin (AlexaFluor488) and counterstained with DAPI. **B)** CT-26 cells were fixed and stained for beta catenin (AlexaFluor488) and counterstained with DAPI. **C)** Quantified nuclear β-catenin in B16F10 cells treated with andrographolide and analogs. **D)** Quantified nuclear β-catenin in CT26 cells treated with andrographolide and analogs.

Qualitatively, several compounds that elicited reductions in cell viability following 24 h of treatment also produced subtle changes in the intracellular distribution of β-catenin. Notably, treatment of B16F10 cells with compounds **3, 4, 6,** and **8** resulted in discrete, intensely fluorescent regions within clusters of cells exhibiting altered morphology, contrasting with the comparatively diffuse fluorescence observed in DMSO-treated cells. These changes were accompanied by differences in cellular density and morphology, suggesting that the redistribution of β-catenin-associated fluorescence may occur alongside treatment-induced changes in cellular organization.

Quantification of nuclear localization of β-catenin by pixel heat analysis is presented in **Figure 9C**. In CT-26 cells, addition of andrographolide and its analogs do not result in any appreciable difference in total amount or subcellular localization of β-catenin, while administration of a CHIR-99021, a GSK-3β inhibitor, expectedly resulted in a significant increase in nuclear localization of β-catenin (p = 0.0026). While B16F10 cells treated with andrographolide and its analogs similarly led to minimal changes in nuclear localization of β-catenin by pixel heat quantification, the administration of CHIR-99021 does not lead to the elevated levels of nuclear β-catenin as was observed in CT-26 cells. This suggests that the earlier observed differences between B16F10 and CT-26 cells in drug sensitivity when cells are exposed to CHIR-99021 may be a result of different sensitivities of the direct target of andrographolide and its C19 analogs towards GSK-3β inhibition.

**Figure 9.**
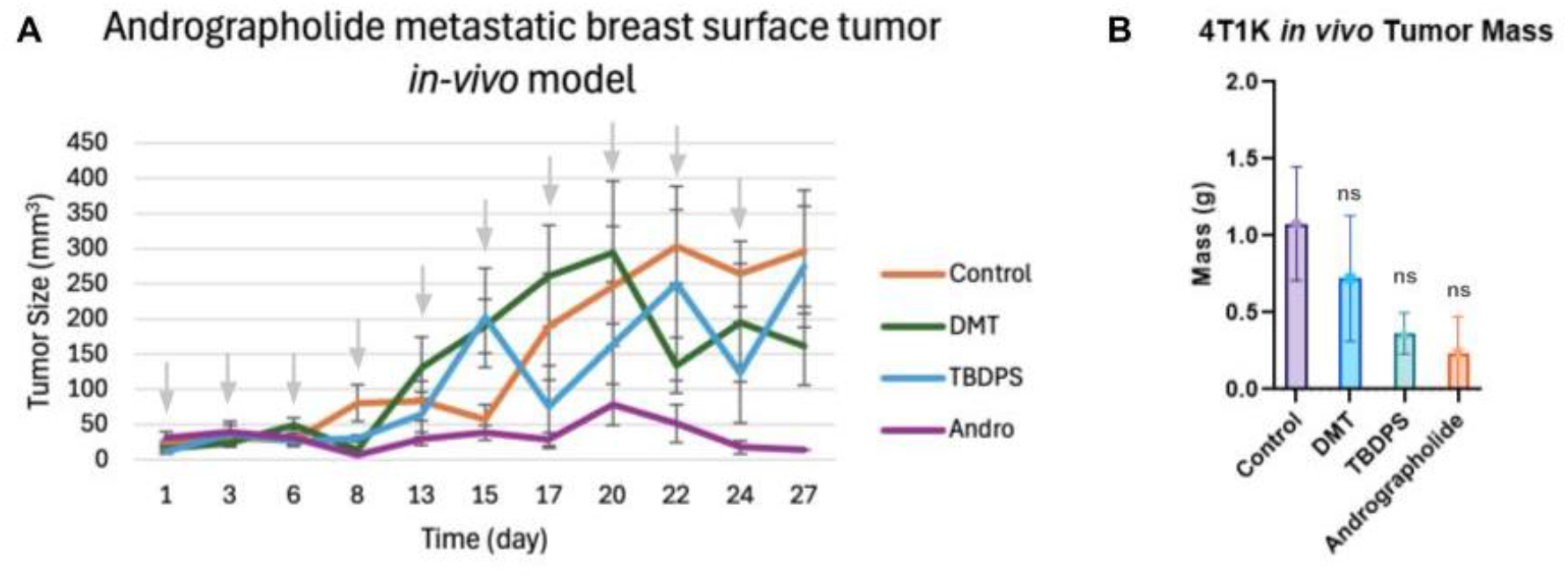
*In vivo* evaluation of lead silyl ether compounds in BALB/c 4T1K murine mammary adenocarcinoma model. **A)** Andrographolide exhibits suppression of tumor growth in a Balb/ 4T1 murine model stably transfected with overexpression of HERV-K. Silyl ether analogs performed marginally better than the DMSO control and inferior to andrographolide in terms of tumor suppression. Arrows represent dosing regimen days. **B)** The overall tumor mass after a 28-day treatment regimen at 0.5 mg/kg. Adjusted p-values are indicated as follows: ns, p≥0.05; *, p<0.05; **, p<0.01; ***, p<0.001.

### 3.8 In Vivo Anti-Tumor Evaluation

Finally, in order to establish *in vivo* anticancer activity of andrographolide and its C19 analogs, we selected two silyl ethers, a *tert*-butyldiphenylsilyl ether (Compound 8) and a dimethylthexylsilyl ether (Compound 9) to be compared against andrographolide and a DMSO vehicle control in a BALB/c 4T1K *in vivo* model. Unfortunately, all silyl ethers exhibited solubility limits far below acceptable projected dosing regimens, and were therefore excluded from *in vivo* trials.

Remarkably, we observe potent tumor-suppressive activity in mice treated with the parent compound, andrographolide **1**, with limited antitumor activity in the *tert-*butyldiphenylsilyl ether **8** with respect to untreated mice **(Figure 9A),** with most mice exhibiting fully suppressed tumors by day 27 with this andrographolide dosing regimen. Upon completion of the 28-day dosing regimen, the difference in mean of endpoint tumor mass is observed with andrographolide displaying the most pronounced difference, followed by the *tert-*butyldiphenylsilyl ether. However, both have limited statistical significance (p = 0.09, p = 0.1034 respectively). **(Figure 9B)**. Comparatively, the dimethylthexylsilyl ether exhibited a moderate but statistically insignificant reduction in endpoint tumor mass **(Figure 9B).**

While both *tert-*butyldiphenylsilyl ether and dimethylthexylsilyl ether exhibited comparable, and sometimes improved *in vitro* potency against 4T1/4T1K cells, this loss of *in vivo* potency may be attributed to dose-limiting solubility challenges of the more lipophilic analogs, which limited maximal dosing concentrations, and may have also impacted tumor penetration. Nonetheless, these studies provide valuable insight into the complex role of silyl ethers of andrographolide in modulating both *in vitro* potency but also *in vivo* bioavailability, in comparison to the more water-soluble natural product.

## Discussion

Andrographolide and its analogs have previously demonstrated potent anticancer activity in several human cancer cell models, highlighting the potential therapeutic utility of this family of labdane diterpenoids. We and others have demonstrated lipophilic functionalization at the C19 hydroxyl group with silyl and trityl ether protecting groups as a strategy for augmenting the biological potency of andrographolide and other bioactive natural products. However, the role of andrographolide and its C19 analogs have not been previously reported in murine cancer cell models. In this study, in order to identify structure-activity relationships of andrographolide and its A-ring analogs in 4T1, CT26, Neuro2a, and B16F10 murine cancer cell lines, we evaluated the antiproliferative activity of C19 silyl and trityl ether analogs of andrographolide at multiple timepoints. From this, we observed potent micromolar-level antiproliferative activity across all four murine cancer cell lines, with unique potency in CT-26 murine colorectal cancer cells.

Next, to identify differences in an *in vitro* model for bioavailability, we performed 2-hour washout experiments, in which cells were only briefly exposed to each compound prior to being displaced with undrugged media. From this, we observed a precipitous loss of *in vitro* potency across both trityl and silyl ether analogs, with slight differences observed in CT26 versus B16F10 cell lines. Further, we generally found that trityl ethers of andrographolide exhibit a much greater loss in potency upon two hour drug washout, suggesting that C19 trityl ethers exhibit slower intracellular uptake and lower bioavailability compared to C19 silyl ether analogs. The expected increase in lipophilicity, coupled with lower aqueous solubility, was quantified computationally, and the increased lipophilicity of C19 functionalized analogs is hypothesized to be a major driver in differences in murine cancer antiproliferative potency and bioavailability of C19 analogs. Further flow cytometry-based experiments demonstrated that silyl and trityl ethers of andrographolide generally do not alter distribution of cell cycle phases in CT26 cells, but significantly differ in their propensity to elicit pro-apoptotic activity, wherein silyl ether analogs 6-13 generally exhibited greater FITC-Annexin V surface staining.

Finally, we sought to identify the role of Wnt1/β-catenin signaling and potential activity of GSK-3β, which are putative targets of andrographolide C19 analogs reported in human cancer cell lines. We found cell line-selective responsive behavior to CHIR-99021, a small molecule GSK-3β kinase inhibitor, wherein the antiproliferative activity of andrographolide and its analogs in B16F10 and 4T1 cancer cell lines were largely unaffected by GSK-3β co-inhibition, while all compounds exhibited a precipitous loss in activity in CT26 cells when co-delivered with CHIR-99021. These findings suggest that CT26 cell lines exhibit a unique GSK-3β sensitivity that is distinct from the mechanisms of action of these compounds in B16F10 and 4T1 cells.

Altogether, we report the biological activity of a library of C19 trityl ether and silyl ether analogs of andrographolide in a panel of murine cancer cell lines. While these compounds exhibit broad-spectrum micromolar-level antiproliferative activity across B16F10, CT26, Neuro2a, and 4T1 cells, further analysis by flow cytometry and CHIR-99021 co-delivery reveal nuanced differences in the precise mode of action through which these compounds exert their anticancer effects. Consequently, the results reported here are consistent with a model that not all putative mechanisms of anticancer activity in human *in vitro* models directly translate to murine models.

## Author contributions

R.R., R.L., A.P., A.X.Y., S.V., and E.N. synthesized the compounds used in this study. R.R., R.L., A.P., A.X.Y., S.V., L.L., L.C., A.H., C.C.M., and E.N. performed cell viability and washout experiments. R.L., A.P., A.X.Y., L.C., A.H., C.C.M. performed apoptosis flow cytometry experiments. L.L. performed *in silico* ADME modeling. H.S. performed cell cycle flow cytometry experiments. A.Y., L.C., and S.V. performed immunofluorescence imaging experiments. R.R., R.L., A.P., A.X.Y., S.V., L.L., L.C. A.Y., H.S., A.H., C.C.M., G.J., F.W.J., and E.N. performed *in vivo* experiments. F.W.J. generated HERV-K env stably transfected cell lines. G.J., F.W.J., and E.N. supervised and conceived of the study. All authors contributed to and approve of the final version of this manuscript.

## Acknowledgements

The authors gratefully acknowledge Tribioscience Inc. for generous support of bioreagents used in this study.

## Abbreviations

4T1K: HERV-K overexpressing 4T1 cells
ADME: Absorption, distribution, metabolism, excretion
Andro: Andrographolide
B16F10K: HERV-K overexpressing B16F10 cells
B16F10: murine melanoma cells
CHIR-99021: a small molecule GSK-3β inhibitor
cLogP: Calculated logarithm of the partition coefficient
Cl Bu (Cl Butyl): 4-chlorobutyldimethyl ether
Cl Trityl: 2-chlorotrityl ether
DMSO: dimethyl sulfoxide
DMT: dimethylthexylsilyl ether
DPM: diphenylmethylsilyl ether
EMT: epithelial-mesenchymal transition
env: Envelope protein
FITC: Fluorescein isothiocyanate
HERV-K: Human endogenous retrovirus type K
KRAS: Kirsten rat sarcoma virus oncogene homologue
MSAB: Methyl 3-{[(4-methylphenyl)sulfonyl]amino}benzoate
MTT: (3-(4,5-Dimethylthiazol-2-yl)-2,5-Diphenyltetrazolium Bromide)
N2A: Neuro2A neuroblastoma cells
NMR: Nuclear magnetic resonance
TBDPS: *tert-*butyldiphenylsilyl ether
TBS: *tert-*butyldimethylsilyl ether
TES: triethylsilyl ether
TPS: triphenylsilyl ether
tPSA: Topological Polar Surface Area
Trityl: trityl ether

